# Cis-Regulatory Differences Explaining Evolved Levels of Endometrial Invasibility in Eutherian Mammals

**DOI:** 10.1101/2020.09.04.283366

**Authors:** Yasir Suhail, Jamie D. Maziarz, Anasuya Dighe, Gunter Wagner, Kshitiz

## Abstract

Eutherian (placental) mammals exhibit great differences in the degree of placental invasion into the maternal endometrium, with humans being on the most invasive end. Previously, we have shown that these differences in invasiveness is largely controlled by the stromal fibroblasts of the maternal endometrium, with secondary effect on stroma of other tissues resulting in correlated differences in cancer malignancy. Here, we present a statistical investigation of the second dogma linking the phenotypic and transcriptional differences to the genomic changes across species, revealing the regulatory genomic sequence differences underlying these inter-species differences. We show that gain or loss of specific transcription factor binding site sequences are connected to the inter-species gene-expression differences in a statistically significant manner, with a particularly larger effect on stromal genes related to invasibility. We also uncover transcriptional factors differentially regulating genes related to pro- and anti- invasible property of stroma. This work extends the understanding of inter-species differences in stromal invasion to the causal genomic sequence differences paving new avenues to target stromal characteristics to regulate placental, or cancer invasion.

## Introduction

Placentation during pregnancy presents a remarkable phenotypic diversity between different eutherian (placental) mammals^1^. This variation is often attributed to the fact that placentation involves the interaction of genetically distinct maternal and fetal tissues ^2–4^. The maternal-fetal interface during placentation is classified anatomically by the depth of invasion into the endometrium as epitheliochorial (e.g. cattle, dugong), endotheliochorial (e.g. dogs, cats and many carnivores), and hemochorial (e.g. humans and other primates, and rodents) ^5–7^. Even among the species with invasive hemochorial placentation, there is large variation in the extent to which placental trophoblasts invade and interact with the maternal tissue ^8,9^. Invasiveness during pregnancy is also correlated with the rate of malignancy among mammals, with significantly lower rate of melanoma metastasis observed among cows, and horses compared to carnivores and humans ^10^.

For long, it was posited that the increased stromal invasion among hemochorial placentation (including in humans) is due to higher invasive capability of placental trophoblasts ^11,12^. Genomically informed phylogeny construction showed that the epitheliochorial placentation has evolved more recently from the ancestral hemochorial placentation, which is present in the stem lineage of eutherian mammals ^6^. We have hypothesized, and experimentally validated that the evolution of epitheliochorial placentation is in part a stromal characteristic, which has resulted in the stromal fibroblasts to attain increased resistance to invasion from placental trophoblasts ^13^. This hypothesis, termed the Evolved Levels of Invasibility (ELI) posits that stromal invasibility itself is an evolved phenotype, resulting in higher resistance to trophoblast invasion within bovines (and other species with epitheliochorial placentation). Notably, its secondary manifestation in other stromal tissues also limit cancer dissemination, suggesting a heritable, genomic basis for the establishment of the phenotype ^13,14^. In this paper, we explore the extent to which genomic changes explain the variation in stromal invasibility in a larger cohort of mammalian species.

Specifically, we characterized the gain or loss of transcription factor binding sites (TFBS) in the promoter region of genes to explain the variation in gene expression between these species. We found that differences in TFBS number could explain many times higher fraction of variance in gene expression for ELI correlated genes compared to other genes, providing further confidence that stromal invasibility is a selected, and inherited phenotype. We also found transcription factors which antagonistically act as potential enhancers of pro-invasive genes, and repressors of anti-invasive stromal genes across eutherians. Because the ELI predicted stromal resistance is secondarily manifested in other stromal tissues, our current work charts a specific inherited genomic basis for the acquired stromal resistance to placental invasion, as well as cancer malignancy.

## Results

### Identifying Gene Expression Signatures Correlated with Placental Invasibility

The ELI hypothesis states that bovine stromal fibroblasts, both in endometrium and skin, have evolved to be more resistant to trophoblast and cancer cell invasion than their human counterparts ^13^ (**Figure 1A**). To explore the evolutionary history of genes that have changed along with non-invasive epitheliochorial placentation, we expanded the number of eutherian species in our transcriptomic analysis. We isolated and cultured the stromal fibroblasts from endometrial tissues of rabbit, guinea pig, rat, cat, horse, humans, sheep, dog, and cow. This selection of species comprise of two eutherian clades, the Euarchontoglires (primates, rodents and others), and Laurasiatheria (ungulates, carnivores and others), together forming the clade of Boroeutheria^5^. This balanced phylogeny contained species with both invasive placentation (human, rabbit, rat, and guinea pig), and less-invasive endotheliochorial or epitheliochorial placentation (cat, dog, horse, sheep, and cow) (**Figure 1B**).

**Figure 1.**
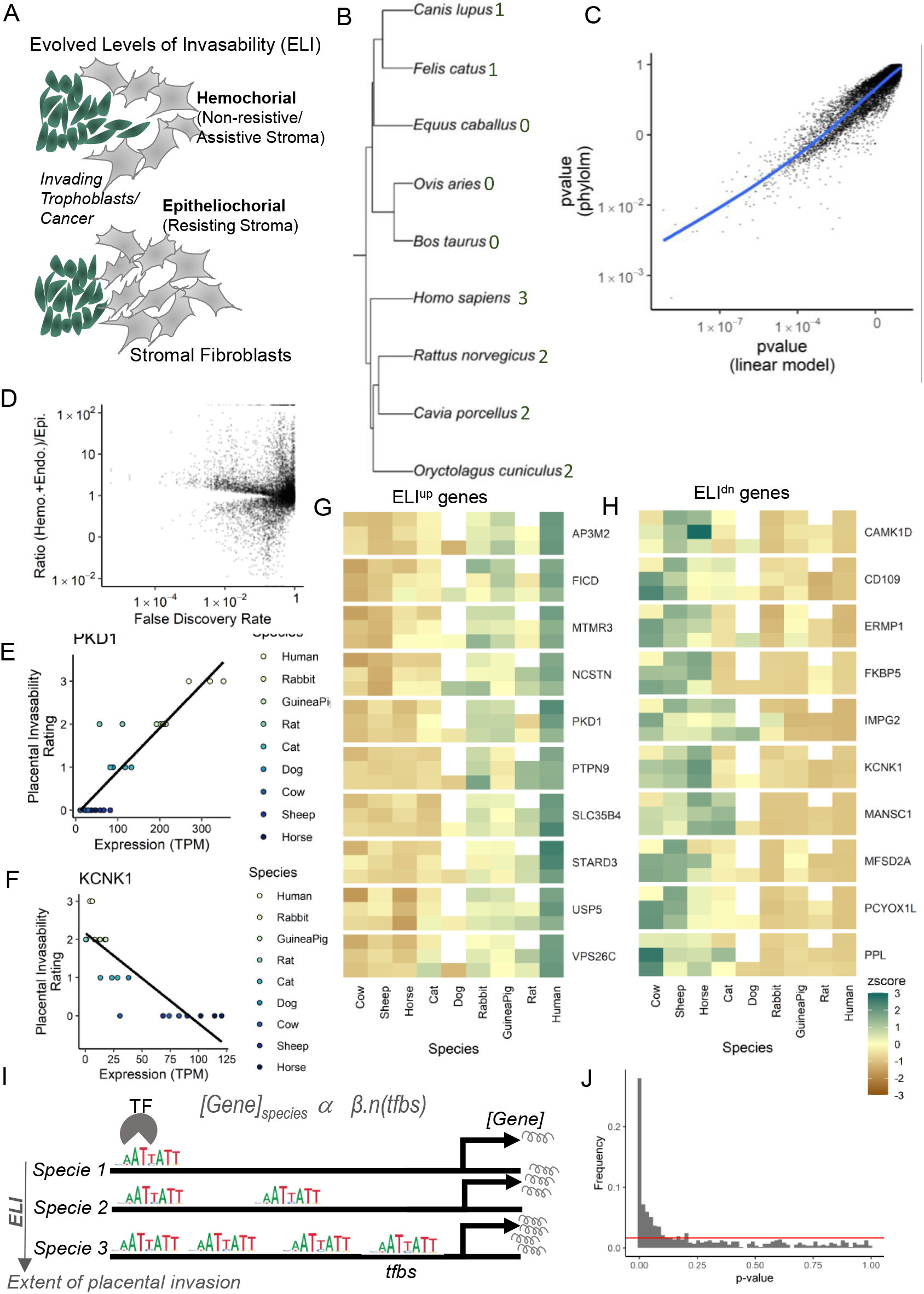
Finding genes correlated with the Evolved Levels of Invasibility (ELI) of the endometrial stroma across eutherian mammals. (A) Schematic explaining the ELI principle, showing that among the epitheliochorial placental mammals, the stroma has evolved to resist trophoblast invasion with secondary manifestation in other tissues resulting in decreased cancer malignancy. (B) Phylogenetic hierarchical representation of selected eutherian species whose endometrial stromal fibroblasts (ESF) were analyzed in the manuscript; The tree is balanced for invasive and less-invasive placental species. Numbers denote the extent of placental invasion ranked from 0 to 3. (C) Statistical significance of the expression of individual genes for correlation with invasibility among different species, evaluated using a simple linear model (x-axis) and a phylogenetic linear model (y-axis). Larger p-values under the phylogenetic linear model, but similar ordering of the genes under the two methods is visible. (D) Individual genes for adjusted p-value (FDR, x-axis) for correlation to placental invasibility against the ratio of the mean expression of the same gene in hemochorial and endotheliochorial species against non-invasive epitheliochorial species. (E) Illustration of the linear fit test used to probe whether the gene expression of a particular gene (in this case, PKD1) in ESFs across species is correlated with the invasibility. (F) Illustration of a gene whose expression is anti-correlated with invasibility. (G and H) Heatmaps of expression across species for the genes most correlated with the invasibility phenotype across species, either positively (ELI^up^ genes), and negatively (ELI^dn^ genes). (I) Schematic showing the gain and loss of binding sites in the promoter regions of a given gene in different species, and its effect on gene expression; b is a coefficient estimating the extent of effect gain of a binding site has on gene expression. (J) The distribution of p-values for individual transcription factor binding site motifs, whose occurrence is tested for an effect on inter-species gene expression differences. The over-representation of low p-values beyond the distribution expected by chance (red line) is evidence of a real effect.

We then created a rough invasibility index based on the anatomical presentation of the fetal-maternal interface in these species^5,15^. Specifically, species with epitheliochorial placentation were given a rank of 0 (cow, sheep, horse), carnivores with endotheliochorial placentation were ranked 1 (cat and dog), rodents with hemochorial placentation ranked as 2 (rabbit, guinea pig and rats), while humans, which along with primates have a particularly invasive hemochorial placentation were ranked 3 (**Figure 1B**). Our objective was to identify genes that correlate with the placental invasibility phenotype. We statistically tested the correlation (or anti-correlation) of gene expression changes in all the species with the placental invasion phenotype with both a naïve linear, and a phylogenetic linear model (see Methods) (**Figure 1C, Supplementary Table 1**). The phylogenetic linear model corrects for a correlation that arise due to shared descent of gene expression and placental invisibility phenotype^16^. As expected, the phylogenetic linear model showed less significant p-values than the naïve model. Nevertheless, the same genes show high correlations in either linear model (**Figure 1C**). Therefore, either model can be used to prioritize candidate genes correlated with invasibility. Notably, we found that overall more genes have stromal expression positively correlated to invasive placentation, rather than negatively-correlated, suggesting that the recent evolution of epitheliochorial placentation is associated with an overall downregulation of genes affecting invasibility (**Figure 1D**).

**Figure 1E** shows an example of a gene with higher expression correlated with increased placental invasiveness (defined as ELI^up^ gene), while **Figure 1H** shows an example of a gene whose expression is anti-correlated to invasibility (defined as ELI^dn^ gene). The ELI^up^ genes are purportedly pro-invasible as their stromal expression increases along with increase in placental invasion, while ELI^dn^ genes are purportedly anti-invasible, as they increase in expression with reduced stromal invasion. **Figures 1G-H** show the top ten genes ranked in order of their positive and negative regression respectively, of their expression and the extent of invasion in each species. We selected a balanced set of 200 genes (all had FDR < 0.05), with 100 most statistically significant genes each, that were correlated (ELI^up^) or anti-correlated (ELI^dn^) with the invasive phenotype. This gene set, termed ELI gene set, is a rank ordered set of genes which show consistent differences across species from the least invasive to the most invasive placentation. This gene set was used to identify candidate transcription factors that regulate stromal invasibility by correlating transcription factor binding site abundance near ELI genes with invasibility phenotype across eutherians.

### Linear Model Relating Transcription Factor Binding Site Abundance to Gene Expression

We used a linear regression model to estimate the extent to which cis-regulatory transcription factor binding site numbers could statistically explain gene expression differences across species. Using gene expression quantified as transcripts per million (TPM), of 8639 genes with 1 to 1 orthology, we first calculated their square root TPM values to decrease the influence of extremely high expression values. We then subtracted the gene-wise mean across all species from the square-root of the gene expression. This scaled expression measure was fit to a linear model with the number of transcription factor binding sites in promoter regions as the predictor variable (**Figure 1I**). Statistical significance and the size of the regression coefficient was recorded for each binding site motif.

Testing for 572 transcription factors for which consensus binding site motifs are available in the JASPAR database^17^, we identified 187 TFs (for a threshold of FDR < .05) which could explain a significant portion of the inter-species differences in gene expression in endometrial stromal cells, as calculated across all genes (**Figure 1J**). A large fraction of TFs statistically influence inter-species gene expression variation indicating that the gene expression profile of endometrial stromal cells has evolved, in part, through changes in cis-regulatory elements (**Figure 1J**).

### Identification of cis-Regulatory Elements Related to Inter-Species Variation in Stromal Gene Expression

We found that β, the regression coefficient describing the extent to which TFBS abundance could explain variation in gene expression, was positive for most TFs (**Figure 2A**). A positive β indicates the gain of TFBS is correlated with increased expression of the target gene, suggesting that most TFs acted as positive regulators of gene expression (i.e., activating TFs) among endometrial stromal fibroblasts (ESFs). The range of value of β was similar for positive and negative regression coefficients (**Figure 2A**). We found that for many TFs, |β| was above 0.025 for the whole transcriptome, with high significance (FDR < 10^−20^) (**Figure 2B, Supplementary Table 2**).

**Figure 2.**
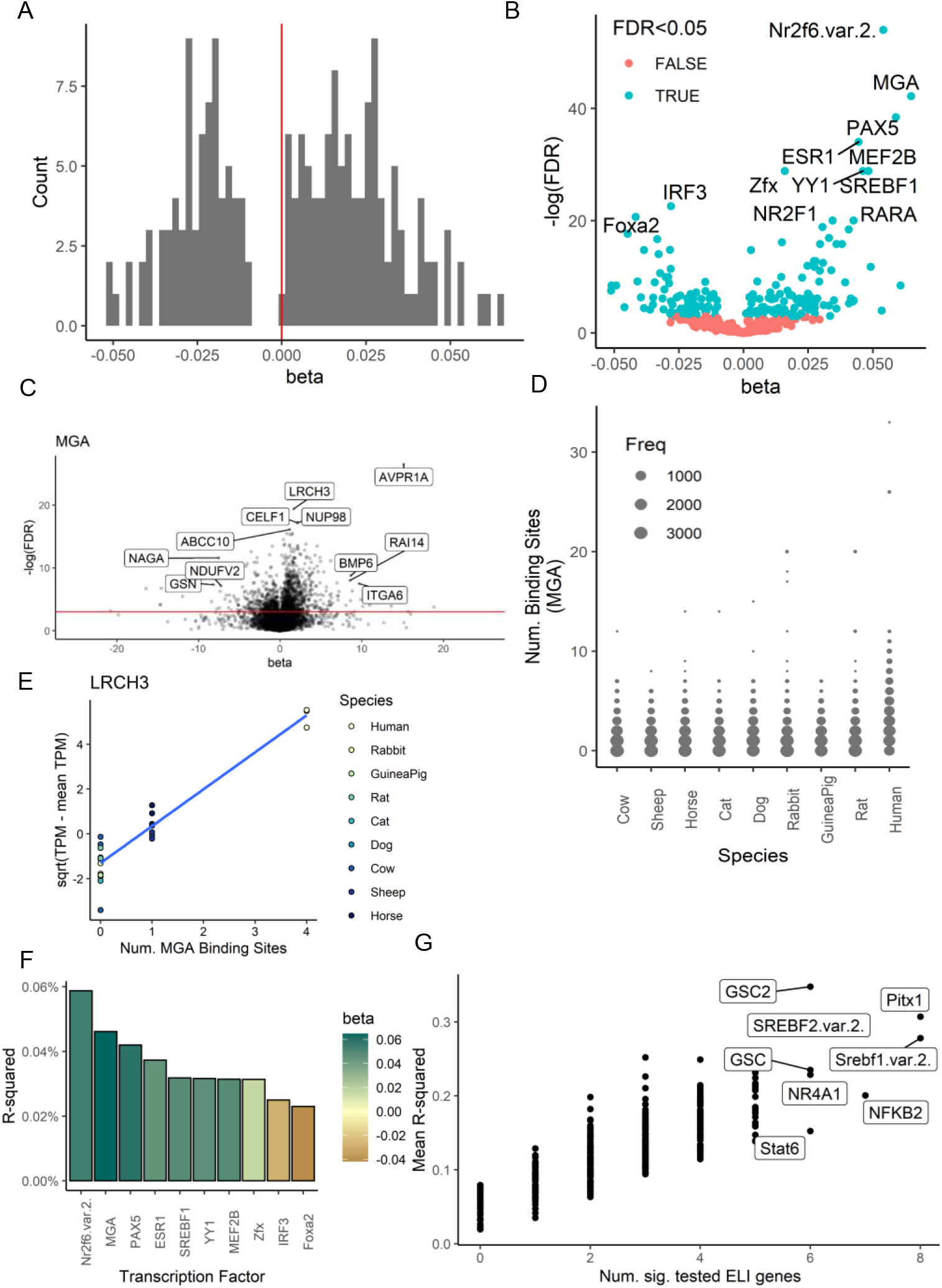
Global analysis of the genomic basis of the modulation of gene expression in endometrial stromal fibroblasts in differential placental invasion among eutherians. (A) Histogram for the effect size (beta) found for different transcription factor binding site (TFBS) motifs that were found to be statistically significant (FDR < .05). Beta represents the amount the scaled (square root of TPM) expression changes with one binding site, on average. Positive beta implies an enhancer effect, while negative beta implies a repressor effect on gene expression of downstream genes. (B) Transcription factors whose putative binding sites affect the variation in expression across selected eutherian species. Each point represents a single transcription factor binding motif. The statistical significance is plotted in terms of the FDR on the y-axis, while the effect size beta (increase or decrease in expression) for additional binding sites is plotted on the x-axis; Statistically significant motifs are labeled green. (C) Illustration of the effect of downstream effect of MGA binding sites on the expression of individual genes. Here, each point represents a gene whose expression was independently tested for a linear relationship to the number of binding MGA binding sites. The gene-specific effect size (beta) and significance (FDR) are plotted on the x- and y-axis respectively. (D) Distribution of the number of binding site motifs matched in the promoter regions of genes in different species, shown here for the MGA binding site motif. (E) Illustration of the effect of MGA binding sites on the expression of LRCH3, across different species listed in the order of placental invasiveness. (F) The R-squared (fraction of variance explained) for the most statistically significant transcription factors affecting global ESF gene expression across species. The colors correspond to the effect size, with blue for an overall increase of gene expression with the particular TF binding in its promoter region, and orange for an overall decreasing effect for additional binding. (G) Transcription factor binding sites whose occurrence is related to the expression of genes experimentally tested^13^ for their role in modulating invasibility. Each point represents a binding site sequence motif, with the mean variance explained on the y-axis and the number of genes with p-value < .05 for the correlation on the x-axis.

Averaged β over all genes did not reflect the large effect a single transcription factor may have on the expression of individual genes (**Figure 2B-C**). As an example, for many genes the regression coefficient |β| for the transcription factor MGA could be 20 times higher than the β averaged over all genes, demonstrating specificity for particular target genes (**Figure 2C**). The inter-species distribution of the number of binding sites for the same transcription factor for all genes showed enrichment for specific TFBS was high for a small number of genes among hemochorial species (**Figure 2D**). As an example Figure 2E shows the expression of the target gene LRCH3, as a function of the number of binding sites for MGA (**Figure 2E**).

We then identified the top transcription factors with the highest R^2^ values (**Figure 2F**). These TFs consisted of various nuclear receptors including NR2F6, a steroid receptor which is also a repressor of IL17, the Estrogen Receptor ESR1, as well as SREBF1, the sterol regulatory element binding transcription factor 1 ^18^. NR2F6 co-regulates and inhibits thyroid receptor^19^, and Oxytocin Receptor (OXTR)^20,21^, while ESR1 directly regulates OXTR expression ^22^, and is considered essential for uterine receptivity in various ruminants^23,24^. In addition, among the top candidates an important factor with a negative β value was FOXA2, the Forkhead Box A2, an essential transcription factor for fertility and uterine function^25,26^. In addition, other top transcription factors with high β values were Max Dimerization Protein MGA, a negative regulator of MYC pathway and implicated in endometrial cancers^27,28^, PAX5 involved in gene regulation in many tissues.

Previously we experimentally demonstrated that human stromal fibroblasts could be rendered more resistant to trophoblast invasion by modifying gene expression to match that of bovine stromal fibroblasts^13^. We tested whether inter-species variation in these experimentally validated ELI genes could be explained by cis-regulatory evolutionary changes (**Figure 2G**). Indeed we found that among 10 experimentally validated genes (ACVR1, BMP4, CD44, DDR2, LPIN1, MGAT5, NCOA3, STARD7, TGFB1, WNT5B), many TFs contributed to their inter-species expression differences. These included homeobox transcription factors like PITX1, and GSC, sterol binding factors like SREBF1, SREBF2, and NR4A1, and nuclear signal transducers like NFKB2, and STAT6. Notably, the experimentally validated genes were positively-correlated with the invasibility phenotype, and accordingly the regulating TFs also had positive β values.

### TFBS Statistically Explain Expression Variation of Genes Correlated with Stromal Invasibility

Our transcriptomic analysis had revealed many genes whose expression is correlated with the placental invasion phenotype, i.e. the ELI^up^ and ELI^dn^ gene sets (**Figures 1C, G-H**). The ELI framework posits that, within eutherians, resistance to placental invasion has emerged by changes in the stromal fibroblasts ^13^. Having shown in the previous section that inter-species variation in gene expression is correlated with TFBS abundance, we focus here on the regulation of the ELI gene set (**Figure 3A**). To account for the possible systemic inter-species variation in the number of binding sites of different transcription factors, we chose the top 100 significant genes each from both ELI-correlated (purportedly pro-invasible, ELI^up^), and ELI-anti-correlated (purportedly anti-invasible, ELI^dn^) gene sets.

**Figure 3.**
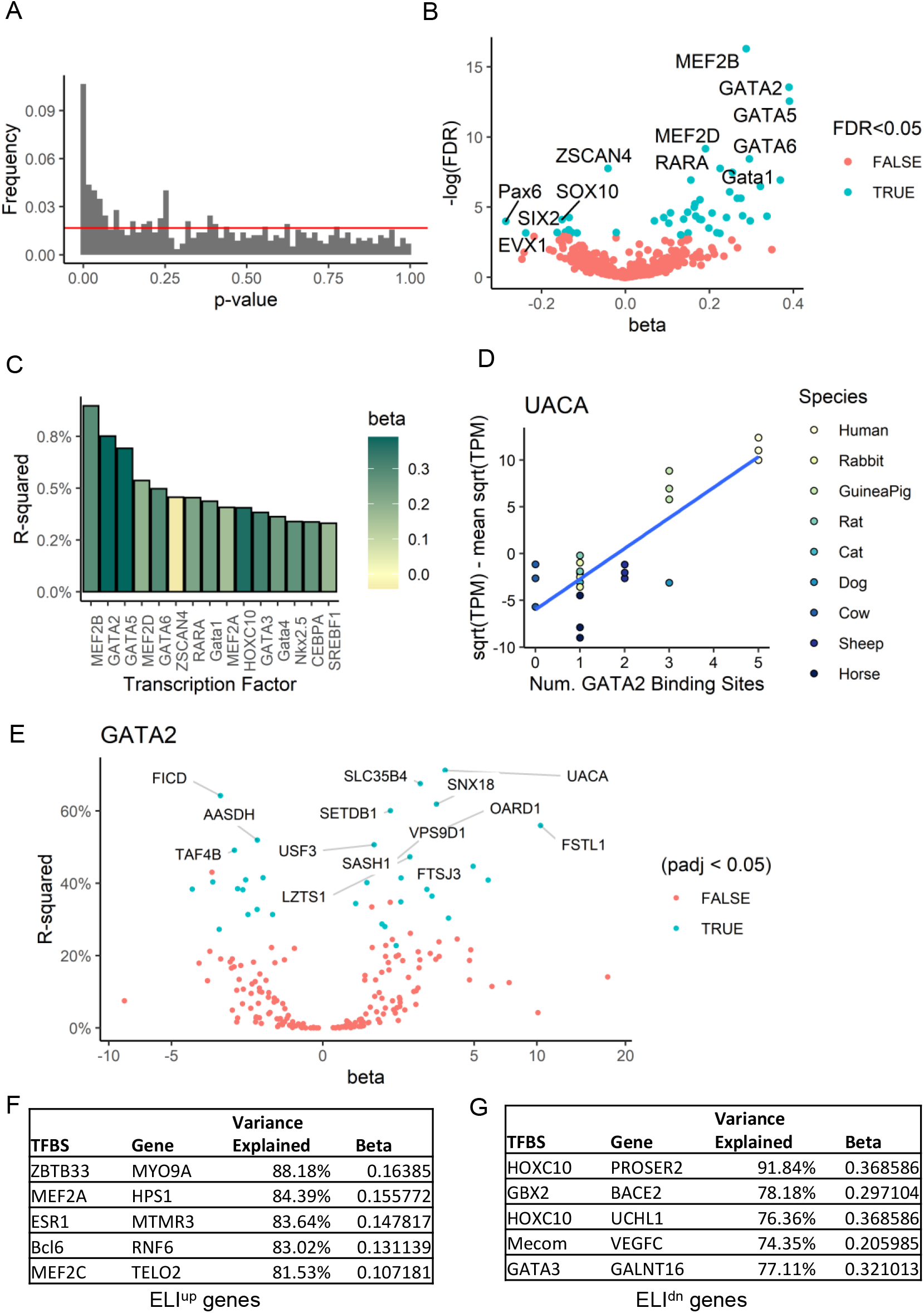
Genomic basis for the regulation of genes correlated with stromal invasibility among eutherian species. (A) The distribution of p-values for the effect of individual transcription factor binding site motifs, tested for the effect on expression of ELI-related genes., with the red line representing the distribution expected by chance. (B) The transcription factors whose binding sites affect the expression of genes purportedly involved in the invasibility of the endometrium (the ELI-related genes). Statistical significance (y-axis, FDR) with the effect size (x-axis, beta). (C) The fraction of variance (R-squared) of the expression across genes explained only by the occurrence of a particular binding motif aggregated over a balanced set of the most significant 100 ELI^up^ and 100 ELI^dn^ genes, shown here for the most significant transcription factor binding motifs. The colors of the bars represent the effect size, green being higher. (D) Illustration of the linear fit used to calculate the statistical significance and effect size for an affected gene, UACA; Shown are the changes in gene expression across species with increasing binding sites for the transcription factor GATA2; Species are ordered on the basis of their extent of placental invasion. (E) Specific genes likely to be regulated by GATA2. The fraction of variance explained (y-axis), and the effect size (x-axis, beta) are calculated for a linear model relating the (scaled) gene expression across species for a single gene to the number of GATA2 binding site motifs in the promoter regions among the different species. (F) ELI-correlated (ELI^up^) and (G) ELI anti-correlated (ELI^dn^) genes whose variance is most explained by the number of specific binding site sequences. Beta is the common effect size found for the binding site sequence found for all ELI genes. For individual transcription factor-gene pair, a large extent of inter-species variance in the direction (or against the direction) of placental invasiveness could be explained by the cis-regulatory TFBS elements.

Modeling specifically for this balanced ELI-specific set of 200 genes, we found that TFBS could statistically explain a much higher fraction of gene expression differences than for all 1-1 orthologous genes (**Figure 3A-B**). For many transcription factors, both the absolute β values, as well as the fraction of variance explained (**Figures 2C, 3C**), were substantially higher than for the whole gene set, with most TFs exhibiting a positive β value. Many of these TFs were related to the myosin enhancer family of transcription factors, with all major MEF TFs, e.g. MEF2A-D, showing significant effects. In addition, we found that GATA transcription factors were highly enriched among the TFs showing significant effects. These included GATA1 and GATA2 which are classically referred to as hemotopoeitic TFs^29^, and GATA5 and GATA6^30^, which are classically referred to as cardiac specific TFs. Crucially, GATA1 and GATA2 are key TFs upregulated in cancer stromal fibroblasts, and induce increased cancer dissemination^31^. All 4 GATA factors are also important in trophoblast development^32^, with GATA2 acting as a key TF regulating decidualization^14,33^. These data highlight genomically encoded transcriptional regulation of decidual genes in species with increased endometrium-trophoblast interaction. Notably, we found relatively few TFs with significant negative β values, but they included key developmental TFs like Pax6, SOX10, and SIX2.

Surprisingly, we found that the transcription factors displaying the highest statistical significance showed R^2^ value 50 times higher for similar calculations aggregated over all genes (**Figures 2C, 3B-C**). Individual genes likely to be involved in regulating placental invasibility (such as UACA) can be seen to be affected by differences in the number of binding sites of particular TFs (such as GATA2) (**Figure 3D**). Further, as for the whole transcriptome, an aggregated R^2^ value for a transcription factor over all ELI genes masked the substantial effect of TFBS in explaining the substantial variance in gene expression for individual downstream genes (**Figure 3E**). As an example, binding sites for GATA2 could explain more than 50% of the inter-species variance for many genes. Interestingly GATA2 is acting either as an enhancer, or a repressor of gene expression (**Figure 3E**). Top TF-gene pairs for TFs which act either as enhancers (β>0), or repressors (β<0) showed that for many ELI genes, a substantial extent of the inter-species transcriptional difference is explained by changes in cis-regulatory binding sites (**Figure 3F-G, Supplementary Table 3**). Overall, these data strongly suggest that hard-coded cis-regulatory genomic changes can significantly explain changes in gene expression associated with increased stromal vulnerability, and stromal resistance to placental invasion.

### Evolution of Increased Resistance to Placental Invasion is Associated with Distinct Patterns of Genomic Regulation

Phylogenetic analysis has settled the debate over the evolution of non-invasive epitheliochorial placentation in eutherians, which is now considered to have evolved after an ancestral invasive hemochorial placentation^6^. However, anatomical reports suggest that among primates, the placentation has continued to evolve into a very aggressively invading form^8,34^, suggesting continued evolution among invasive placentation. We asked whether cis-regulatory genomic changes could specifically explain gain or loss of the phenotype of placental invasibility. Towards this objective, we tested if ELI genes that correlate with increased (or decreased) placental invasion are differently influenced by TFBS.

We used top 100 genes which were the most positively correlated (ELI^up^ genes), and the top 100 genes most anti-correlated (ELI^dn^ genes) with the invasive placental phenotype within the stroma (FDR < 0.05 for all selected genes). Separate treatment of either gene-sets in our model could be influenced by systematic inter-species differences in TFBS frequencies, because the ranked order of ELI^up^ (or ELI^dn^) genes is correlated with the placental invasion phenotype, and hence the species, by definition. To avoid this artefact, we normalized the frequencies of each TFBS across the species. We found that a higher number of TFs showed gene expression enhancing effects for ELI^dn^ genes, while for ELI^up^ genes, relatively more TFs showed repressive effects on gene expression (**Figure 4A**).

**Figure 4.**
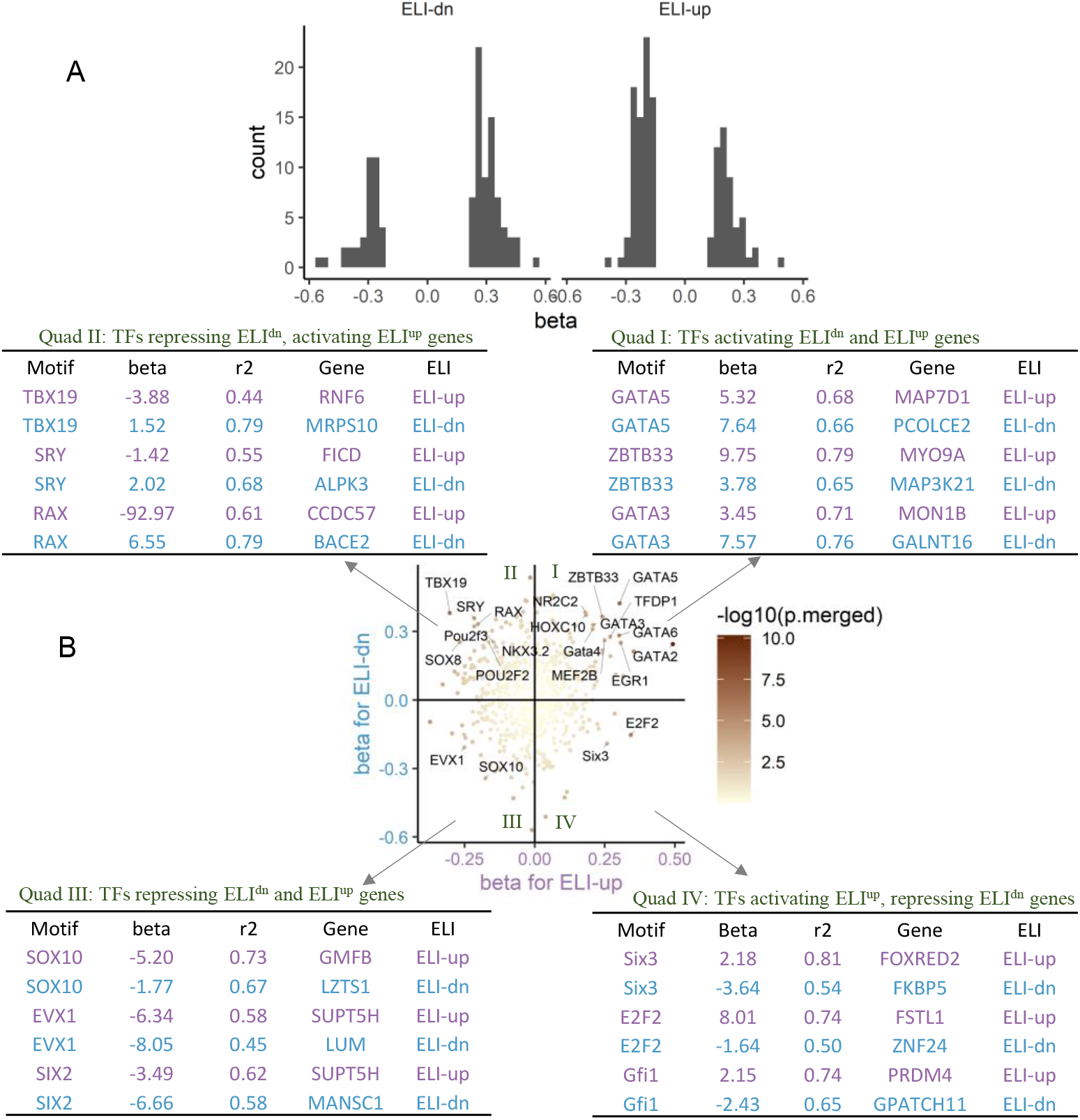
Differences in the TFBS explained regulation of genes correlated (ELI^up^), or anti-correlated (ELI^dn^) with placental invasiveness in endometrial stromal fibroblasts. (A) Histograms of the effect sizes (beta) of different binding sites, when tested on only ELI^dn^ and ELI^up^ genes. Only effect sizes for TFBSs that are statistically significant (p-value > 0) in the respective genes (ELI^dn^ or ELI^up^) (B) Scatter plot shows the effect sizes (beta) of individual TFs for ELI^dn^ (y-axis) and ELI^up^ (x-axis) genes. Names of some selected TFBS’s are labeled. Selected genes with a high fraction of variance explained by TFBS motif occurrences are shown as examples in the four tables next to each quadrant.

For each TF, we analyzed the effect size for either gene sets (ELI^up^ and ELI^dn^) after normalizing for TFBS occurrence across species (**Figure 4B**). We found that while few TFs act as either enhancers or repressors for both ELI^up^ and ELI^dn^ genes (quadrants I, and III), there were also TFs which acted as antagonistic regulators for ELI^up^ and ELI^dn^ genes (quadrants II, and IV). Tables show examples of TFs which regulate a given ELI^up^, and ELI^dn^ gene either as an activator or a repressor. Indeed, for many genes, a large fraction of their inter-species variation is explained by a single TF. The statistical significance of the regulation by each TF for individual ELI genes, as well as for the two sets of 100 ELI^up^ and ELI^dn^ genes are listed in Supplementary Table 3.

Quadrant I lists TFs activating both ELI^up^ and ELI^dn^ genes, and includes many GATA factors, and MEF2 factors. All GATA factors, and MEF2 factors are key TFs regulating human and mouse ESF decidualization. We speculate that these TFs may regulate resistive genes in ESFs of epitheliochorial placentation, and also genes which are present in the more resistive decidualized fibroblasts in hemochorial species. Other TFs also suggest a link to decidualization and preparation for pregnancy including estrogen responsive factor, HOXC10^35^, and early growth response gene 1 (EGR1)^36^. EGR1 is induced by placenta growth factor and is an inducer of hypoxia inducible factor 1 (HIF1α) necessary for early implantation^37^. Quadrant III shows TFs which acted as repressors for both ELI^up^ and ELI^dn^ genes.

In contrast, quadrant II shows TFs which act as repressors for ELI^up^ genes, and activators for ELI^dn^ genes. Since ELI^dn^ genes are increased in species with epitheliochorial placentation, these TFs may act as important regulators promoting stromal resistance, by both activating resistive genes, as well as downregulating genes which potentially promote invasion. These TFs include POU2F2/3, SRY, Nkx3.2, and TBX19. Quadrant IV lists TFs, including SIX3, and E2F2 acting as antagonistic repressors of pro-invasibility ELI^up^ and activators of anti-invasibility ELI^dn^ genes.

Overall, these analyses provide a map of genomically encoded changes in stromal gene expression that have accompanied placental invasion across species. Presence of antagonistic transcriptional regulators, which act oppositely on pro-invasibility ELI^up^ and anti-invasibility ELI^dn^ genes suggest that the vastly different phenotypic outcomes related to placentation are achieved partially by shared mechanisms. For example, through direct genomic binding, TBX19, acts as an activator for MRPS10, and a repressor for ANGEL1 associated with increased placental invasion. These antagonistic transcription factors also present attractive targets to limit invasive processes by gene knockdowns.

## Discussion

The maternal fetal interface presents a highly diverse physiology among placental mammals with large inter-species differences in the extent of placental invasion into the endometrium, even though the central function of this interface of nutrient transfer from the mother to the fetus is broadly conserved^38^. Phylogenetic analysis has proven that the non-invasive epitheliochorial placentation is a more recent innovation than the invasive hemochorial placentation^6,39,40^. This has led to the rejection of the long held hypothesis that the hemochorial placentation, also present in humans, had evolved recently. We have advanced a hypothesis, termed “ELI” (Evolved Levels of Invasibility) stating that evolution of non-invasive epitheliochorial placentation was driven by changes in the endometrial stromal genotype, i.e. less invasive placentation is caused by higher stromal resistance to invasion^13^. Experimental validation proved that human stromal invasibility can be modulated by regulation of gene expression similar to that in epitheliochorial mammals. Here, we used an ordinal metric of placental invasiveness to identify candidate genes which are correlated with invasibility among eutherians using transcriptomic data of ESFs from 8 species, balanced for invasive and non-invasive placentation. Both a naïve linear model, and a phylogenetic adjustment to the model identified similar genes, which we termed as ELI^up^ and ELI^dn^ genes according to their correlation with placental invasibility phenotype.

Because the invasibility phenotype varies between species, one expects that genomic differences would inform inter-species transcriptomic differences in ESF between those presenting invasive and non-invasive placentation. Further support of a genomic basis for the phenotype of stromal invasibility was its secondary manifestation in other tissues, e.g. skin, wherein epitheliochorial species present lower rate of malignancy^10,41^, which we have also experimentally validated^13^. In this work, we explored an important aspect of hard-coded genomic regulation, the abundance of transcription factor binding sites in promoter regions, and asked how well these explain the transcriptomic differences between eutherian species. We found that TFBS for individual transcription factors could, to a limited extent, explain inter-species variaion in gene expression.

Indeed, TFBS explained a much larger fraction of inter-species variation in ELI^up^ and ELI^dn^ genes. For individual TF/target gene pairs, the cis-regulatory elements themselves could explain a large portion of expression changes along the trajectory of the evolution of reduced placental invasion. TFs explaining large ELI-correlated transcriptomic differences belonged to myocyte enhancer binding factor 2 (MEF2) families, and GATA-binding factor families. The binding sites for these TFs were also more abundant in species with stronger placental invasion, e.g. rodents and humans, with a concomitant effect on increased downstream gene expression. We speculate that this may be due to a more tissue specific phenomenon, decidualization, which occurs primarily in hemochorial species. MEF2 proteins, in particular MEF2B and MEF2D, are central players in muscle cell differentiation^42,43^, cardiac hypertrophy, fibrosis^44^, and stroma induced epithelial to mesenchymal transition of cancer^45^. Similarly, GATA factors are also important for the endometrial decidualization program. GATA2 in fact, is a central factor in endothelial cell differentiation^46^ and decidual differentiation in humans and mice^14,33^. The stroma of hemochorial species may be more prepared to interact with invasive trophoblasts, and among primates, also be participants in their endothelial transformation. All GATA factors found to have significant influence on invasibility in our study are also crucial for trophoblast development in hemochorial species^32^. Indeed, GATA2 has been reported to regulate genes associated with decidualization, and decidua-trophoblast interactions^47^, including placental lactogen synthesis^48,49^, proliferin^47^, and prolactin-like protein-A^50^. GATA5/6 are also important for cardiogenesis, similar to MEF2A and MEF2B. Whether these transcriptional regulators act as pro-invasive factors will need to be experimentally tested. Interestingly, GATA2 gene is alternatively spliced in non-placental species, with the −3.9kb regulatory element of the gene absent in non-eutherians^32^. Epitheliochorial placentation among ruminants is a more recent derivation than the ancestral hemochorial placentation. Therefore it is worth noting that it not the gain of binding sites for MEF2, GATAs and other TFs, but in fact the loss of these binding sites is found in species with reduced stromal invasion.

In addition to the TFs which acted as enhancers for pro-invasibility genes, we found many TFs which acted antagonistically on ELI^up^ and ELI^dn^ genes. Many of these antagonistic TFs (Figure 4D, quadrant II) acted as enhancers for anti-invasibility (ELI^dn^) genes, while also acting as repressors for the pro-invasibility (ELI^up^) genes. What do these TFs tell us about the evolution of epitheliochorial placentation? Because evolutionarily, epitheliochorial placentation is more recent, our analysis suggests that both, ELI^dn^ and ELI^up^ genes have recruited these transcription factor binding sites, but their effects on ELI^dn^ and ELI^up^ genes are opposite. Indeed, we also found examples of antagonism wherein certain TFs (e.g. E2F2 in Figure 4D, quadrant IV) act as repressors of ELI^dn^ genes, while activating ELI^up^ genes. Here, the target genes have lost binding sites for these TFs with the evolution of epitheliochorial placentation. We underline that while the effect of these antagonistic TFs is opposite upon the expression of ELI^up^ and ELI^dn^ genes, the effect on the phenotype of invasibility is consistent. TFs in quadrant II are expected to be anti-invasibility and those in quadrant IV are expected to be pro-invasibility through their regulatory effect on ELI^up^ and ELI^dn^ genes within stroma. Our analysis presents attractive candidates to test for their effect on stromal invasibility. This analysis also provides a cue to understand the transcriptional mechanisms which have accompanied recent increase in placental invasiveness among the primates.

The ELI hypothesis posits that the stromal compartments of ruminants are particularly resistant to collective cell invasion, by trophoblast or by malignant cancer cells (e.g. skin). Here we have identified a number of candidate transcription factor genes that likely regulate invasibility of endometrial stromal cells, and potentially other stromal fibroblasts regulating cancer dissemination. Understanding the molecular functions of these transcription factors in stromal cell – trophoblast interaction will reveal the mechanisms that enable stromal cells to resist invasion by both, placental as well as cancer cells.

## Methods

Endometrial stromal fibroblasts were isolated and sequenced from 9 species and gene expressions calculated for each sample. A set of 8,639 orthologous gene sets with members in each of the species were identified for analysis. The expression of these genes was calculated in transcripts per million (TPM).

### Finding genes related to the modulation of placental invasibility

We obtained the gene expression *e_g,s_* of gene *g* in species s in transcripts per million. For each species, we also rated the degree of placental invasibility *r_s_* ∈ Error! Bookmark not defined.. In order to find the stromal genes likely involved in modulating the degree of invasibility, we fit the linear model

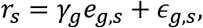

where *ϵ_g,s_* is the residual. This is fitted independently for each gene, and positive *γ_g_* represent genes that are up-regulated in species with more invasive placenta, while negative *γ_g_* indicates the opposite. The statistical significance is calculated for the null hypothesis of *γ_g_* = 0. The p-values for all genes are corrected for multiple testing using the false discovery rate method. In addition, this linear model was also fitted using a phylogenetic linear model, with an assumption of Brownian motion processes producing both the traits (gene expression and species invasibility). The phylogenetic linear model was fitted using the phylolm package^16^.

### Genomic explanation of transcriptomic variation

For fitting the variation in gene expression between species to the cis-regulatory elements, we first calculated a scaled version of the gene expression by first calculating the square roots and then subtracting the mean expression of each gene across all species. Using the gene expression *e_g,s_* for gene *g* in species *s*, we calculate the scaled gene expression as

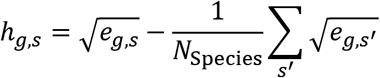

Known eukaryotic transcription factor binding site motifs were downloaded from the JASPAR database^17^. Genomic sequence FASTA files were downloaded from the Ensembl database^51^. We identified genomic regions 5kb upstream to 1kb downstream of the translation start site of each gene as purported promoter regions. We searched for matches to each known binding site motif in each of these promoter regions using the FIMO package^52^ using default parameters. The number of statistically significant matches (p-value < 10^−4^) for each binding site motif in each gene’s promoter regions were assembled in a matrix. We represent the number of matches to the binding site motif of transcription factor *t* in the promoter region of gene *g* in species s as *n_t,g,s_*. Then, independently for each transcription factor binding site motif, a linear model was fit for a set of all genes (or all ELI genes). The linear model related the gene expression between species to the differing number of binding site matches found in the promoter regions of the orthologs of the same gene across all 9 species. The inter-species variation in gene expression was modeled to relate to the occurrences of the transcription factor binding sites as

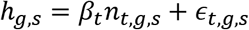

where *ϵ_t,g,s_* is the residual and *β_t_* is the fitted coefficient for a particular transcription factor binding site motif *t*. A model is fitten independently for each motif *t*. Again, a p-value is obtained for the null hypothesis *β_t_* = 0. The coefficient (*β_t_*) of this linear regression model, and its statistical significance (p-value) was recorded for each transcription factor motif.

---

**Supplementary Table 1**: Scoring genes to the Evolved Levels of Invasibility (ELI), in terms of correlation and p-values of gene expression with endometrial invasibility.

**Supplementary Table 2**: Results of statistical testing each transcription factor binding site (TFBS) for their effects on the global inter-species gene expression profile.

**Supplementary Table 3**: Transcription factors found to be regulating pro-invasive (ELI-up) and anti-invasive (ELI-dn) genes.

